# Mitochondrial Metabolism and Calcium Handling in Parkinson’s Disease hiPSC-derived Astrocytes

**DOI:** 10.64898/2026.08.07.743508

**Authors:** Giovanna C. Cavalcante, Camille C. Caldeira da Silva, Éverton L. Vogt, Felipe G. Ravagnani, Victória A. Fulaneto, Patricia de Carvalho Aguiar, Alicia J. Kowaltowski

## Abstract

Parkinson’s disease (PD) is the second most common neurodegenerative disorder worldwide, and mutations in the *LRRK2* and *PRKN* genes are among the most common familial causes of the disease. In neurodegenerative diseases such as PD, disturbances in Ca²⁺ homeostasis and cellular bioenergetics impair the function of neurons and glial cells, contributing to disease progression. These changes are not limited to neurons; mitochondrial dysfunction and disrupted Ca^2+^ homeostasis in astrocytes are increasingly recognized as key contributors to PD, impairing bioenergetics, redox balance, neuroinflammatory responses, and metabolic support essential for dopaminergic neuron survival. In this study, we investigated mitochondrial calcium homeostasis, mitochondrial oxidative phosphorylation, morphology and distribution in human induced pluripotent stem cell (hiPSC)-derived astrocytes with mutations in the PD genes *LRRK2* (G2019S) and *PRKN* (c.155delA; Ex3-4del) and wild-type controls. Intracellular calcium dynamics were assessed using Fura-2 AM. Compared with control astrocytes, *LRRK2*-related PD patient-derived mutant astrocytes exhibited lower intracellular calcium levels, and slower calcium extrusion following stimulation with ATP. Mitochondrial morphology was analyzed using MitoTracker Deep Red, revealing increased mitochondrial fragmentation and redistribution of mitochondria toward the cell periphery in both PD mutant cell types. Because oxidative phosphorylation is tightly regulated by mitochondrial morphology and calcium homeostasis, we next assessed oxygen consumption rates using a continuous metabolic monitoring system (Resipher) and quantified the expression of genes (RT-qPCR) and proteins (capillary electrophoresis-based western detection) involved in mitochondrial calcium transport and bioenergetics. These analyses showed that *PRKN* mutant astrocytes exhibit a more oxidative bioenergetic phenotype than *LRRK2* mutant astrocytes, while both mutant lines displayed altered phosphorylation of mitochondrial morphology regulator DRP1 as well as decreased levels of respiratory complexes relative to control astrocytes. In summary, this study identifies astrocyte-specific mitochondrial dysfunctions and calcium dysregulation as key features of *LRRK2*- and *PRKN*-related pathology, providing new insights into how glial metabolic alterations contribute to neurodegeneration in PD.

## Introduction

Parkinson’s disease (PD) is the second most common neurodegenerative disorder worldwide and is characterized by the progressive loss of dopaminergic neurons, leading to motor symptoms including bradykinesia, tremor, postural instability, and muscle rigidity, as well as non-motor manifestations such as depression, anxiety, cognitive impairment, and dementia (Reich & Savitt, 2019; Jankovic & Tan, 2020; Simon et al., 2020). Among genes associated with familial PD, mutations in *LRRK2* and *PRKN* are common and have been extensively used to investigate disease mechanisms.

The *LRRK2* gene (Leucine Rich Repeat Kinase 2) in the PARK8 locus is an active participant in central biological processes such as autophagy, oxidative phosphorylation, mitochondrial morphology, dynamics and removal through mitophagy (Bonello et al., 2019; Monzel et al., 2023). Several mutations are associated with late-onset Parkinson’s disease (PD), such as G2019S, with symptoms very similar to the sporadic form of the disease (Verma et al., 2017), leading to reduced PINK1/Parkin-dependent mitophagy activity in PD (Bonello et al., 2019).

The *PRKN* gene (parkin RBR E3 ubiquitin protein ligase), also known as *PARK2* or *Parkin*, actively participates in mitophagy (degradation of damaged mitochondria). Mutations in the *PRKN* gene, such as c.255delA, a 1bp frameshift deletion that leads to truncated protein synthesis, are associated with the early development of PD (Abbas et al., 1999; Muñoz et al., 2002; Marder et al., 2010). The main mitophagy pathway is the PINK1/Parkin-dependent pathway, which induces engulfment of target mitochondria with subsequent degradation by lysosomes. When dysregulated, this pathway is related to PD (Cowan et al., 2018; Uoselis et al., 2023). Additionally, Parkin has been shown to be related to calcium homeostasis (Pereira et al., 2023).

Neurodegenerative diseases are universally characterized by alterations in calcium homeostasis and bioenergetics that impair the normal activity of neural cells, as these cells possess an exceptionally high demand for energy and oxygen, and rely heavily on mitochondrial ATP production. Consequently, dysfunctions in mitochondrial energy metabolism directly influence neurodegenerative pathogenesis (Panchal & Tiwari, 2019). Calcium is an important regulator of energy metabolism, with strong signaling effects in the brain. Therefore, imbalance in calcium homeostasis, and particularly mitochondrial calcium homeostasis, can contribute to neurological diseases (Dey et al., 2020). In mitochondria, calcium homeostasis is controlled by influx transporters, such as MCUc (a complex that encompasses subunits MCU, MICU1, MICU2, MICU3, MCUB, MCUR1, and EMRE), and efflux transporters, such as NCLX and TMEM65 (Garbincius et al., 2025).

While the effects of PD on mitochondria and calcium homeostasis in neurons are extensively explored (Glaser et al., 2019; Dey et al., 2020; Boag et al., 2021), much less is known about astrocytes, particularly in human cells, despite the fact that astrocytes are known to be intimately involved in PD (Kam et al., 2020; Kim et al., 2023; Kim et al., 2026). In this study, we investigated calcium homeostasis and mitochondrial metabolism in hiPSC-derived astrocytes with specific PD-related mutations in *LRRK2* (G2019S) and *PRKN* (c.155delA; Ex3-4del) genes.

## Methods

### Cell reprogramming and differentiation

The following PD patients’ skin fibroblasts cell lines were obtained from the Coriell Institute for Medical Research: ND29542 (heterozygous mutant for *LRRK2* G2019S) and ND29543 (compound heterezygous mutant for *PRKN* with c.155delA and Ex3-4del). Wild-type Primary Human Dermal Fibroblasts (HDFa, Gibco™, C0135C) were used as controls. Fibroblasts were cultured in DMEM (Gibco™ #11885084) supplemented with + 10% fetal bovine serum (Gibco™, #12657029) and 0.2% Antibiotic-Antimycotic (Gibco™, #15240112) in a CO_2_ incubator (5% and 37°C).

Based on a previously published protocol (Ravagnani et al., 2023), we reprogrammed and differentiated the human fibroblasts into astrocytes. Fibroblasts were reprogrammed into iPSCs using non-integrative episomal vectors from the ReproRNA™-OKSGM Kit (STEMCELL Technologies, #05930) according to the manufacturer’s instructions. After 28 days, single iPSC colonies were manually collected and transferred to Matrigel^®^-coated plates (Corning^®^, #CLS354227) in Essential 8™ Flex Medium (Gibco™, #A2858501). Clones were passaged every 4-5 days at a split ratio of 1:2 or 1:3 using 0.5 mM EDTA in DPBS (Gibco™, #14190235). For the first 24 hours after each passage, the medium was supplemented with 5 µM ROCK inhibitor Y-27632 (Sigma, #SCM075).

For differentiation into NPCs, iPSC clones at approximately the 5^th^ or 6^th^ passage were cultured for 48 hours in Neurobasal (NB) medium consisting of DMEM-F12 with GlutaMAX™ (Gibco™, #10565018), with 1% B27 (Gibco™, #17504044) 0.5% N2 (Gibco™, #17502048) and 2% Normocin (InvivoGen, #ant-nr-2), which was further supplemented with 1 µM Dorsomorphin (Sigma #P5499) and 10 µM SB431542 (Sigma # 54317). On the third day of the protocol, for embryoid body (EB) formation, the iPSC colonies were scored using a pipette tip, gently detached from the plate using a cell lifter (Corning^®^, #CLS3008), and transferred to 6-well ultra-low-attachment plates (Corning^®^, #3471) on an orbital shaker at 90 rpm inside the incubator. On the third day, the medium was replaced with NBF medium, consisting of NB supplemented with 20 ng/mL each of FGF2 (Gibco™, #PHG026) and EGF (PeproTech^®^, #AF-100-15-500UG). The EBs were kept on the shaker for five additional days, with the medium changed every other day. After this period, the EBs were collected in a Falcon tube, briefly dissociated with Accutase™ (StemCell Technologies, #07920), washed with PBS 1× (Gibco™, #10010023) and centrifuged at 1000 rpm for 3 minutes. After removal of the supernatant, the EBs were resuspended in NBF medium and seeded onto Matrigel^®^-coated plates, with the medium changed every other day until neural rosettes were detected. For differentiation into neural progenitor cells, the neural rosettes were manually collected and seeded onto plates coated with 10 µg/mL poly-L-ornithine (Sigma #P3655) and 2.5 µg/mL laminin (Sigma #L2020 or Gibco™ #23017-015), in NBF medium, which was changed every other day. When the NPCs reached 80 to 90% confluency, residual neural rosettes were removed via vacuum aspiration. The remaining NPCs on the plate were briefly dissociated with 0.5× TrypLE (Gibco™, #12604039) and transferred to new poly-L-ornithine/laminin-coated plates in NBF medium at a 1:2 ratio. The medium was changed every other day until the cells reached 90% confluency, at which point they were passaged at a 1:3 split ratio. To investigate aneuploidies at the fourth passage, all NPCs clones were analyzed by karyotyping. Only normal clones proceeded to differentiate into astrocytes.

For astrocyte differentiation, NPCs at 90% confluency were briefly washed with 1× PBS, supplied with fresh NBF medium and then detached from the plate with a cell lifter. Cells were subsequently dissociated by pipetting and transferred to 6-well ultra-low-attachment plates, on an orbital shaker at 90 rpm inside the incubator. On the following day, neurospheres were observed and the medium was replaced by NBF with 5 µM ROCK inhibitor. After 48 hours, the medium was switched to the AGM™ Astrocyte Growth Medium BulletKit™ (Lonza, #CC-3186). The neurospheres were kept in the shaker for 15 days with the AGM changed every other day. The neurospheres were then collected, seeded onto 10 µg/mL poly-L-ornithine/ 5 µg/mL laminin-coated plates and kept in AGM medium. Astrocytes were passaged every 5-7 days, at a 1:2 or 1:3 ratio. This process was performed for each clone, WT and mutants. Experiments were carried out with one clone per condition. Cell counts were performed using Countess 3 FL (Thermo Fisher Scientific, #AMQAF2000).

### Immunocytochemistry

The astrocytic phenotype was confirmed by evaluating *CD44* and *GFAP* markers in wild-type cells (passage 6) using immunocytochemistry (ICC). Primary antibodies used were CD44 (156-3C11) Mouse Monoclonal Antibody (Cell Signaling, #3570) at 1:200 and Anti-Glial Fibrillary Acidic Protein (GFAP) rabbit polyclonal antibody (Millipore, #AB5801) at 1:300. The secondary antibodies used were Donkey anti-Mouse IgG (H+L) Highly Cross-Adsorbed Secondary Antibody, Alexa Fluor™ 488 (Invitrogen, #A-21202) at 1:400 and Donkey anti-Rabbit IgG (H+L) Highly Cross-Adsorbed Secondary Antibody, Alexa Fluor™ 568 – (Invitrogen, #A10042) at 1:400, respectively. The nucleus was marked using DAPI staining.

### Fluorescence microscopy

For fluorescence-based microscopy experiments, cells were seeded onto 35 mm compartmentalized CELLview glass-bottom dishes (Greiner Bio-One, #627870) pre-coated with poly-L-ornithine and laminin (10 µg/mL and 5 µg/mL, respectively). A total of 10^4^ cells were plated in 500 µL of culture medium per compartment and maintained for 48–72 hours before calcium imaging or mitochondrial morphology analyses. All experiments were performed with 3 to 4 replicates, and images were acquired using a Leica DMi8 inverted fluorescence microscope (Leica Microsystems) operated with LASX Software (Leica Microsystems).

### Intracellular calcium measurements

Fura-2-AM (Invitrogen, #F1221) was resuspended in DMSO to form a 1 mM stock solution. On the day of the experiment, cells plated as described for microscopy, above, were incubated with 5 µM of the probe for 30 minutes at 37°C in 1 mL calcium-free PBS, washed with 500 µL of PBS, then resuspended in 500 µL of PBS with CaCl_2_ (2 mM). A filter wheel installed in the light source permitted Fura-2 fluorescence excitation at 340 nm (Ca^2+^-bound) or 387 nm (Ca^2+^-free), with emission at 510 nm. Baseline cytosolic calcium levels were monitored for the initial 4 minutes. ATP (100 µM) was added as a stimulus for cytosolic calcium increase from organellar release. The total duration of each experiment was 8 minutes. To automate image analysis from each individual cell region of interest (ROI), scripts were developed in the Python programming language 3.9.7 capable of identifying individual cells and mitochondrial location (central versus peripheral) within each cell. The scripts are openly available at: https://github.com/cavalcantegc.

### Mitochondrial morphology assessments

Mitochondrial morphology was assessed by microscopy using the MitoTracker Deep Red FM probe (Invitrogen, #M22426), which accumulates in active mitochondria of living cells. Cells plated for microscopy, as above, were labeled two quadrants at a time, at most, to preserve stability of the probe. The MitoTracker solution was prepared by adding 1 µL of the probe to 199 µL of PBS, from which 5 µL were used in 500 µL, for a final probe concentration of 50 nM. Samples were incubated at 37°C for 20 minutes, followed by washing with 500 µL of PBS and then imaging. The probe excitation wavelength was 644 nm, while the emission wavelength was 665 nm. Captured images were analyzed for morphology using the ImageJ software with the Mitochondria Analyzer plugin. To estimate mitochondrial morphology and (perinuclear or peripheral) location in each cell, a script was developed in Python 3.9.7 programming language using the PyCharm environment (v. 2021.3.2). More information on this script can be found in Cavalcante & Kowaltowski (2026).

### Oxygen consumption rates (OCR)

Oxygen Consumption Rates (OCR) were monitored in real time over 72 hours using the Resipher platform equipped with a 32-sensor lid (Lucid Scientific, #NS32-101A), under standard CO_2_ incubator growth conditions. Astrocytes were seeded at a density of 10-15x10^3^ cells per well in 96-well flat-bottom plates, pre-coated with poly-L-ornithine/laminin, in a final volume of 100 µL of culture medium per well. Each clone was analyzed in triplicate, and cells were allowed to adhere for at least three hours before starting the assay. To minimize edge effects and media evaporation during the 72-hour incubation, all peripheral and unused wells were filled with an equivalent volume of PBS or sterile media. OCR recorded continuously at 37°C and 5% CO_2_. Raw OCR data were background-subtracted using media-only control wells, and the data were normalized by the number of seeded cells. The normalized AUC was measured for the 24-72h range.

### RNA extraction

For RNA extraction, the sample was frozen at 80°C, then thawed at room temperature with 500 µL of TRI Reagent (trizol) (Sigma-Aldrich, #T9424), vortexed, and incubated for 5 minutes. Chloroform was added at a 1:5 v/v ratio to trizol, the sample was vortexed and incubated for 10-15 minutes at room temperature. Samples were centrifuged at 12,000 g for 15 minutes at 4°C, and then a volume of 200 µL of the aqueous phase was carefully transferred to another tube. A 1:1 volume of isopropanol was added, the sample mixed, incubated for 10 minutes at -20°C, and then centrifuged (12,000 g for 15 minutes at 4°C). The supernatant was removed and the pellet washed with 1 mL 75% ethanol. Centrifugation was then performed at 7,500g for 5 minutes at 4°C, the liquid content removed, and the tube left open to dry the pellet, which was then resuspended in 10 µL of water. RNA was quantified using a NanoDrop 1000 spectrophotometer (Thermo Fisher Scientific). For cDNA conversion, the High-Capacity cDNA Reverse Transcription Kit (Thermo Fisher Scientific, #4368814) was used, with 1 µL enzyme, 10 µL buffer, and 9 µL sample for 1 µg of RNA input, using the following thermocycler program: 37°C for 1 hour, 95°C for 5 minutes, and 4°C “hold” to remove from the thermocycler and freeze at - 20°C until qPCR was performed.

### Gene expression

Quantitative real-time PCR (RT-qPCR) was used with human 18S rRNA gene (h18S) as the endogenous normalizer. Primer pairs for qPCR were designed and acquired, as shown in Table 1. Each gene was assessed with 3-6 replicates.

**Table 1.**
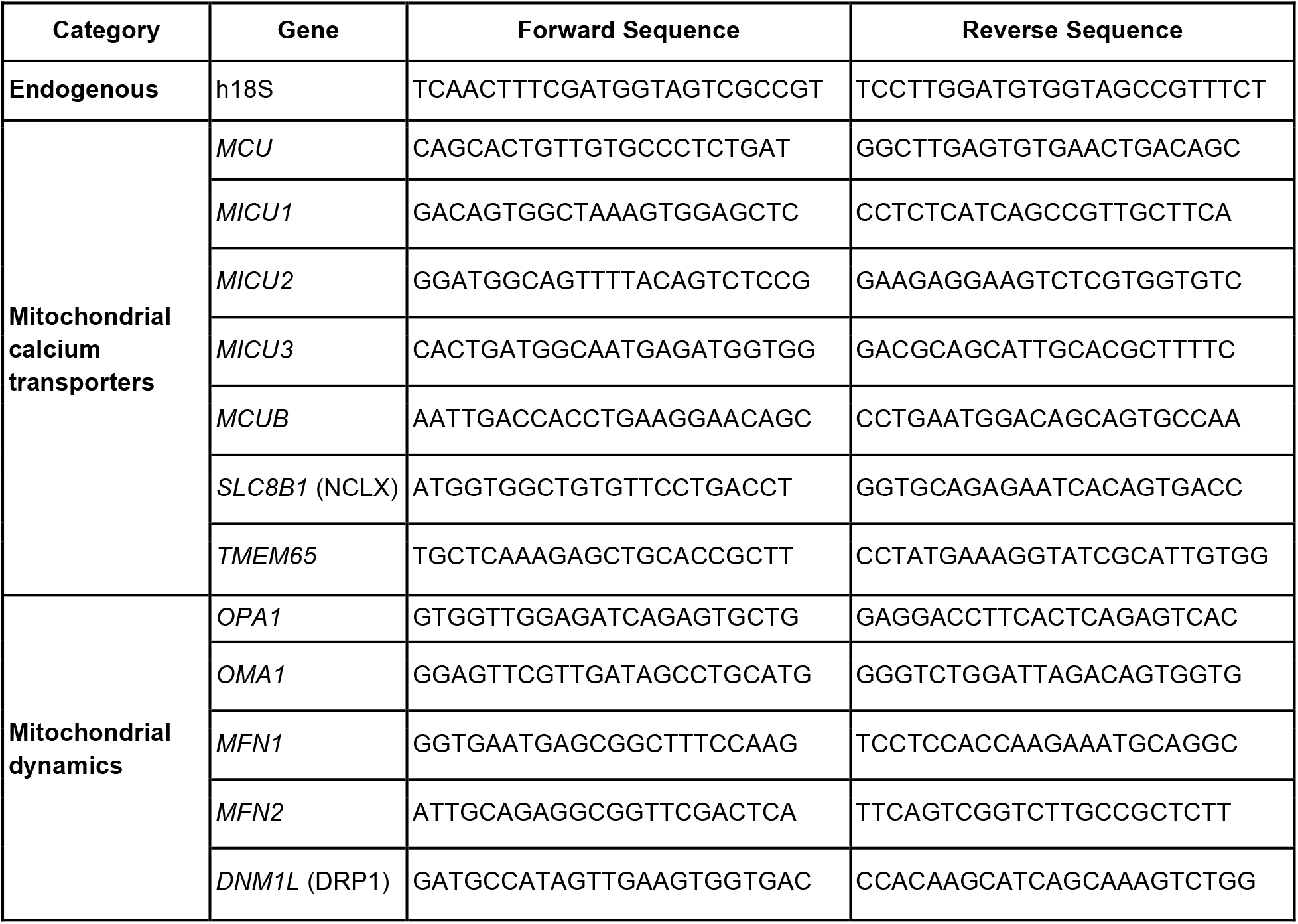
Primer pair sequences used in gene expression assays.

The method used in the real-time equipment (QuantStudio™ 3 Real- Time PCR System, Thermo Fisher Scientific) was the standard obtained in the Diomni™ Design and Analysis 3.0.1 program (Thermo Fisher Scientific). The sample protocol was optimized from the standard 25 µL to a 10 µL total volume, consisting of 5 µL Platinum™ SYBR™ Green qPCR SuperMix-UDG (Invitrogen, #11733046), 0.2 µL Forward Primer, 0.2 µL Reverse Primer, 0.2 µL ROX Reference Dye (part of the kit), 3.4 µL of water, and 1 µL of sample (10 ng/µL cDNA). To confirm astrocytic identity, the expression of the specific markers *GFAP* and *CD44* was quantified via qPCR, providing molecular validation of the hiPSC-derived astrocyte phenotype (results not shown).

### Protein expression

Cell lysates were maintained in RIPA buffer (Sigma-Aldrich, R0278) with protease and phosphatase inhibitors (Thermo Scientific, 78442), and protein quantification were performed using the Qubit Protein BR Assay kit on a Qubit 4 fluorometer (Thermo Fisher Scientific, #Q33239).

Protein expression was assessed using a quantitative capillary electrophoresis-based immunoassay in the Jess Automated Western System (Bio-Techne, #004-650). All assays were performed using 1 µg of each sample, and the chemiluminescence kit for detection (PROT-DM001-KIT for Rabbit or PROT-DM002-KIT for Mouse), following the manufacturer’s instructions. Three to five replicates were performed for each target protein.

Table 2 presents a description of the antibodies used for the investigated proteins and their standardized dilution conditions. Unless otherwise stated, all target protein antibodies were multiplexed with anti-Vinculin antibody for normalization/loading control. Secondary antibodies were used according to the requirements of the primary antibody, being either Rabbit or Mouse, included in the chemiluminescence kits. All data were processed using Compass for Simple Western software.

**Table 2.** List of antibodies used in the protein expression assays.

| Primary Antibody | Catalog Number | Molecular Weight (kDa) | Dilution | Secondary Antibody |
| --- | --- | --- | --- | --- |
| Anti-Vinculin | 4650S | 124 | 1:50 | Rabbit |
| Anti-MCU | 14997S | 30 | 1:100 | Rabbit |
| Anti-MICU1 | 12524S | ~47 | 1:50 | Rabbit |
| Anti-MICU2 | HPA045511 | 50-55 | 1:50 | Rabbit |
| Anti-MICU3 | HPA024771 | ~60 | 1:50 | Rabbit |
| Anti-MCUB | HPA048776 | ~40 | 1:50 | Rabbit |
| Anti-SLC24A6 (NCLX) | 21430-1-AP | 64, ~45 | 1:50 | Rabbit |
| Anti-TMEM65 | ab236861 | 25 | 1:20 | Rabbit |
| Anti-MFN1 | 14739S | 82 | 1:100 | Rabbit |
| Anti-MFN2 | 9482S | 80 | 1:100 | Rabbit |
| Anti-OPA1 | 80471S | 80-100 | 1:100 | Rabbit |
| Anti-OMA1 | 95473S | 37 | 1:100 | Rabbit |
| Anti-DRP1 | 8570S | 78-82 | 1:50 | Rabbit |
| Anti-p-DRP1 (Ser616) | 3455S | 78-82 | 1:50 | Rabbit |
| Anti-Total OXPHOS (R) | 45-8099 | 20, 30, 40, 48, 55 | 1:50 | Mouse |
| Anti-NDUFA9 | ab14713 | 37 | 1:50 | Mouse |
| Anti-SDHA | ab14715 | 70 | 1:50 | Mouse |

### Data analyses

Unless otherwise indicated, all data analyses were performed with the GraphPad Prism v.8.4.2 software or Python v.3.9.7 in the PyCharm v.2021.3.2 environment. Statistical analyses were conducted with Wilcoxon signed-rank test (linked dot plot of Fura-2-AM assays), Kruskal-Wallis with Dunn’s test (Resipher assays), and One-way ANOVA (all other analyses). Grubbs’ test (alpha=0.05) was used to detect outliers. In all cases, statistical significance was considered for P < 0.05.

## Results and Discussion

We began by deriving cell lines from PD patients with specific mutations in *LRRK2* (heterozygous mutant for G2019S) and *PRKN* (heterozygous mutant for c.155delA and Ex3-4del), as well as controls (wild-type for these mutations) by reprogramming skin fibroblasts into iPSCs, and then promoting differentiation into neural progenitor cells and, subsequently, astrocytes (Figure 1 shows a schematic of the workflow; Figure 2 shows representative images of the cells during reprogramming and differentiation). *CD44* and *GFAP* expression were assessed by PCR (not shown) and immunocytochemistry to ensure that astrocyte markers were present in the derived cells (Figure 3).

**Figure 1.**
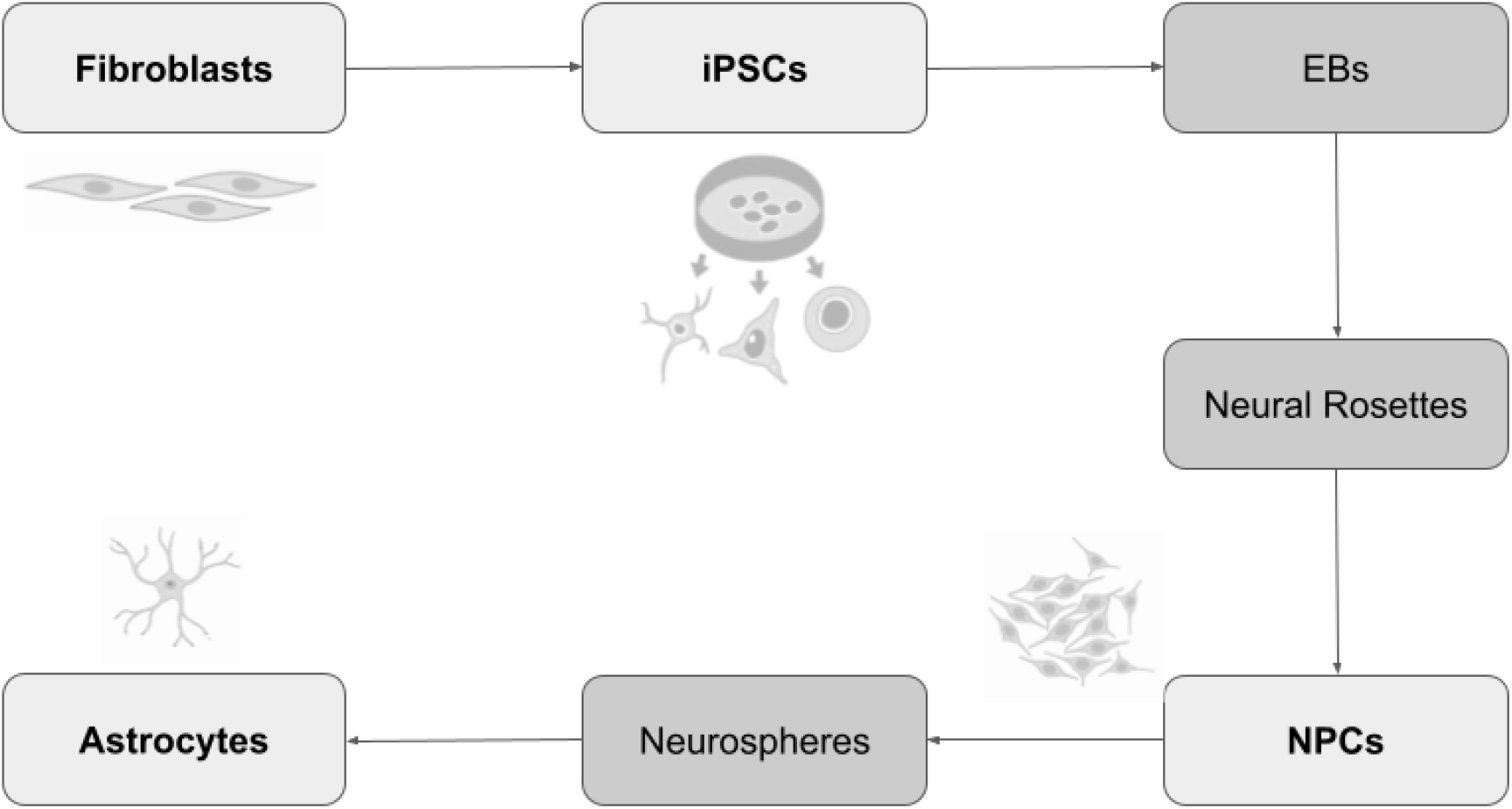
Schematic workflow of fibroblasts reprogramming into induced pluripotent stem cells (iPSCs) and subsequent differentiation into neural progenitor cells (NPCs) and astrocytes. Core cell lines are represented in light gray, while the intermediate stages (embryoid bodies - EBs, neural rosettes, and neurospheres) are represented in dark gray. Created with BioRender.

**Figure 2.**
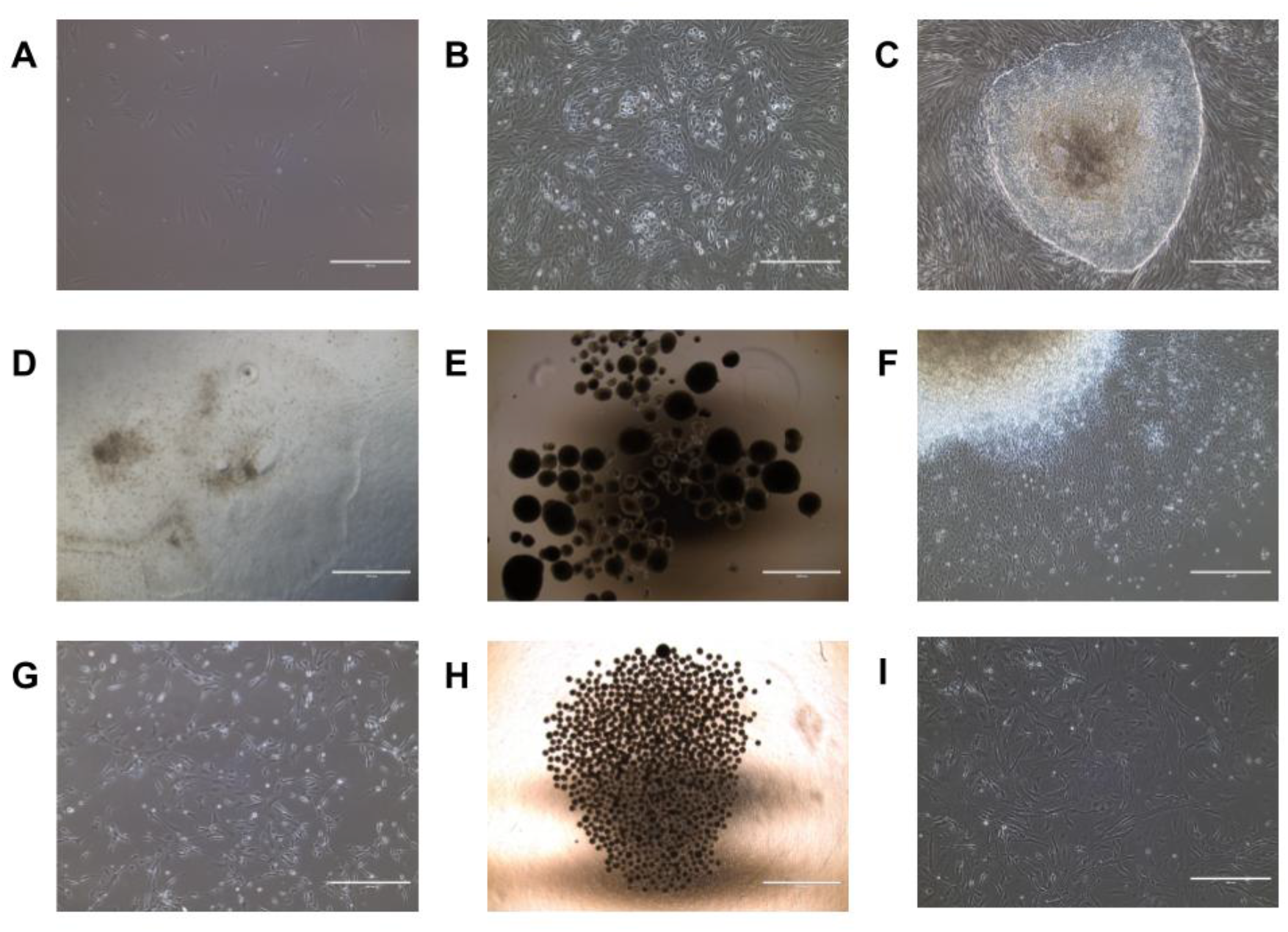
Cell types at each stage of reprogramming and differentiation. Phase-contrast images acquired with an EVOS^®^ XL Imaging System (Thermo Fisher Scientific) with magnifications of 4x (Panels A-C, F, G, I), 10x (Panel D), and 20x (Panels E, H). **A)** Fibroblasts. **B)** Fibroblasts undergoing reprogramming (day 9). **C)** Initial iPSC colonies (day 21). **D)** iPSCs. **E)** Embryoid bodies. **F)** Neural rosettes. **G)** Neural progenitor cells (NPCs). **H)** Neurospheres. **I)** Astrocytes.

**Figure 3.**
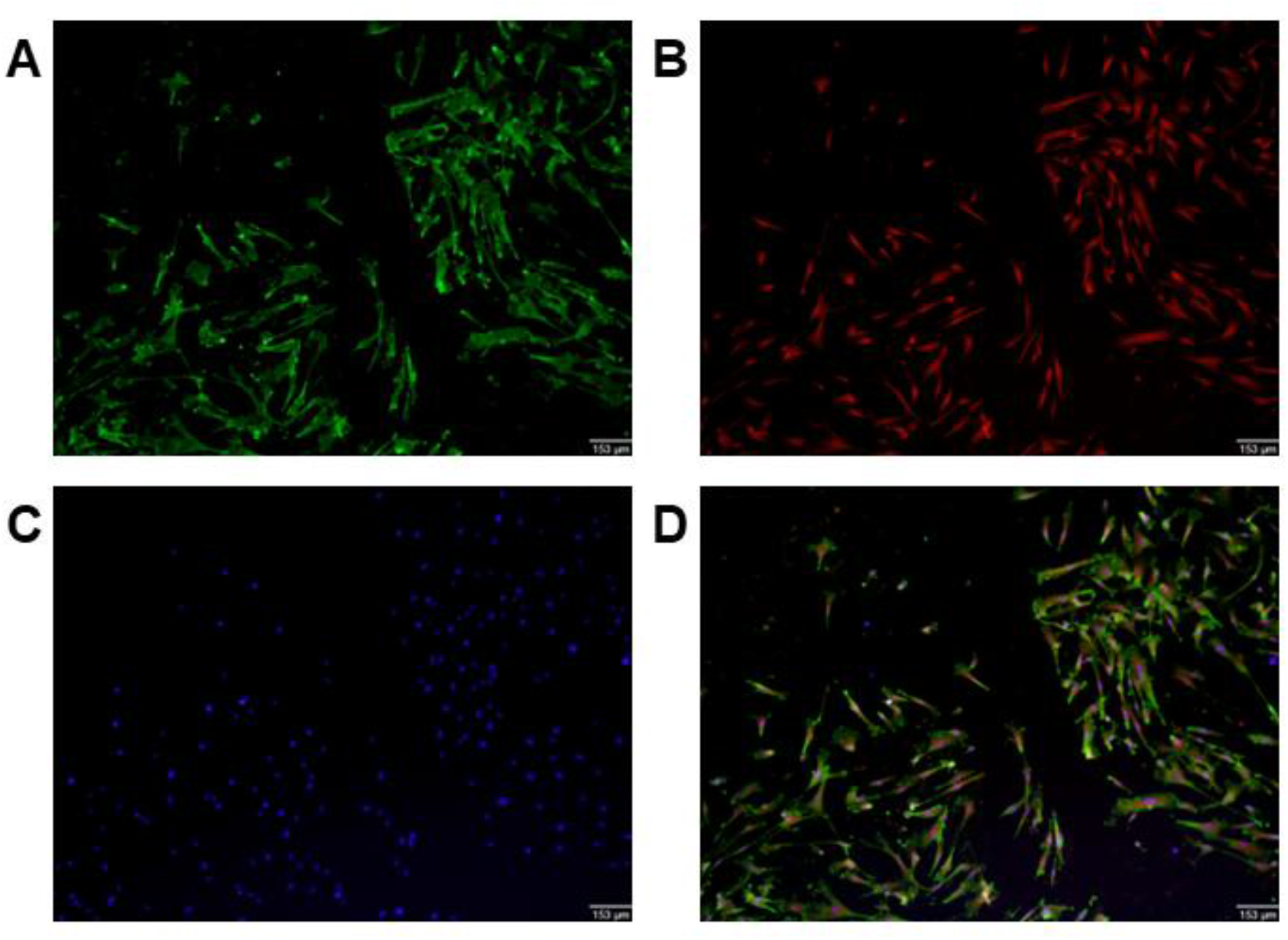
Immunocytochemistry detection of astrocyte markers in wild-type control cells for validation of astrocyte identity using fluorescence microscope Olympus FSX100 with magnification of 4x. **A)** CD44 marker in green. **B)** GFAP marker in red. **C)** Nucleus staining with DAPI in blue. **D)** Merged fluorescence.

Changes in astrocytic Ca^2+^ dynamics have been increasingly recognized as essential for central nervous system function, regulating many functions, including metabolic states and mitochondrial activities (Volterra and Meldolesi, 2005; Veiga et al., 2025; Sanchez-Mico et al., 2026). We measured intracellular Ca^2+^ levels and responses in astrocytes using the ratiometric intracellularly loaded probe Fura-2, a probe which shifts its absorption properties when bound to calcium, so that ratios of the bound to unbound fluorescence peaks (340/380 nm) correspond quantitatively with changes in intracellular Ca^2+^ concentrations. Cells were loaded, and ROI detection was automated, with ratios measured at baseline and after intracellular levels were increased by stimulation with ATP (Figure 4A-D), which binds to purinergic receptors and triggers calcium release from intracellular stores such as the ER into the cytosol (Song et al., 2007). *LRRK2* mutant astrocytes loaded with Fura-2 displayed markedly decreased baseline levels of intracellular calcium compared to WT astrocytes, while *PRKN* mutant astrocytes presented similar baseline levels compared to the controls (Figure 4 C, D, and E). This difference was maintained after the injection of ATP (Figure 4C and F). The decay time (in seconds) of intracellular calcium levels to baseline after ATP addition was also estimated and was significantly increased primarily in *LRRK2* mutant cells (Figure 4G). Overall, patient-derived PD mutant astrocytes displayed abnormal calcium homeostasis, a finding compatible with previous findings in *LRRK2*-mutant astrocytes and neurons (Schwab & Ebert, 2015; Ludtmann et al., 2019), but different from results in *PRKN*-mutant neurons and neuroblastoma cells (Sandebring et al., 2009; Yokota et al., 2023).

**Figure 4.**
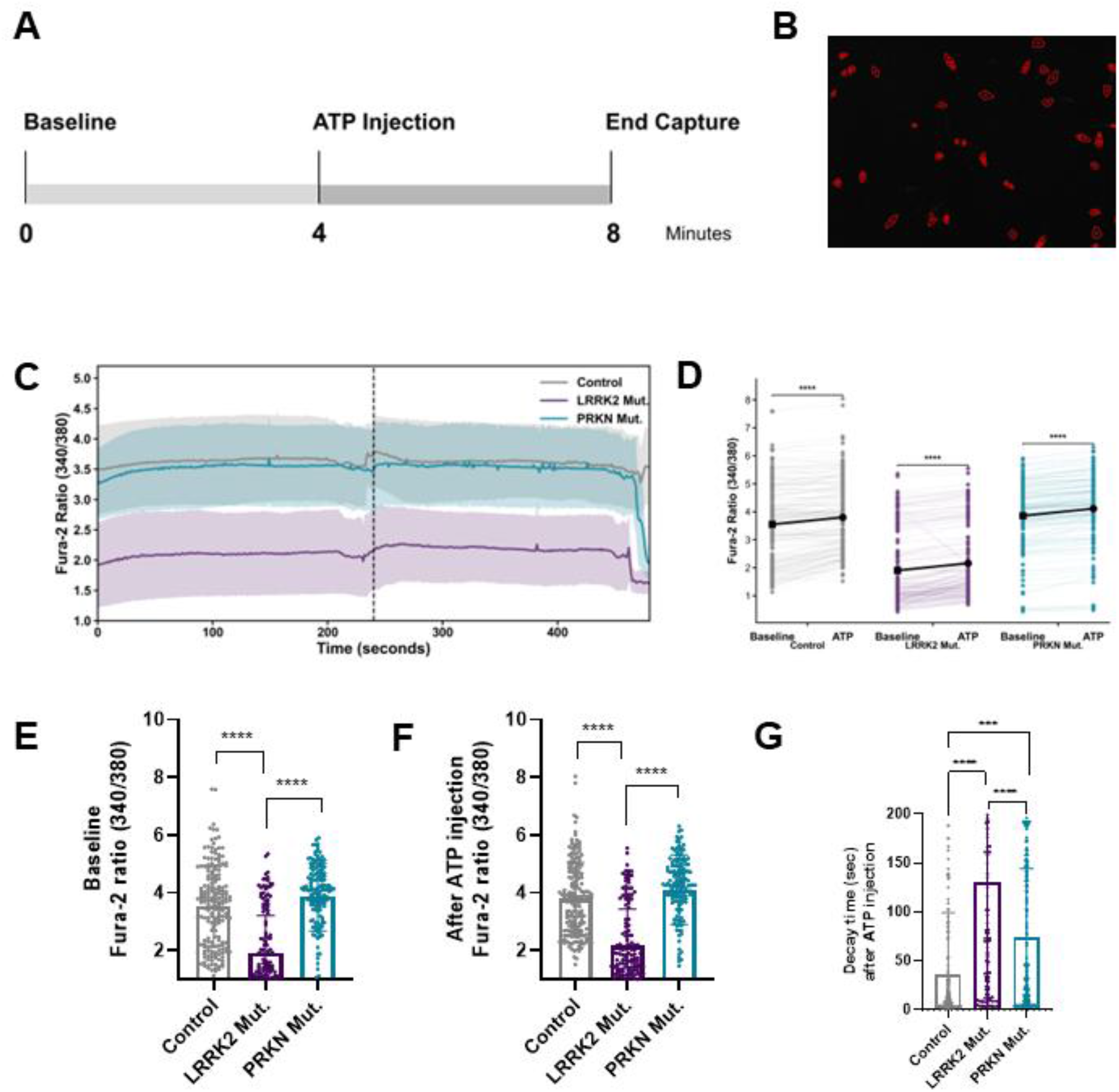
Intracellular calcium homeostasis is altered in PD astrocytes. **A)** Timeline of the process in minutes with each intervention step marked. **B)** Representative automated ROI acquisition (in red). **C)** Mean traces of ROI calcium levels from the time-lapses obtained. The dotted line marks the ATP addition (100 µM). **D)** Linked dot plot showing the Fura-2 ratio for each ROI before and after ATP injection. **E)** Fura-2-AM ratio (340/380) at baseline. **F)** Fura-2-AM ratio (340/380) after the ATP injection. **G)** Time (in seconds) of decay of levels after ATP injection. Graphs represent Means ± SD. Symbols show individual cell data. Wilcoxon signed-rank test (Panel C) ***P_adj_ < 0.0001. One-way ANOVA (Panels E-G) ***P_adj_ = 0.0002; ****P_adj_ < 0.0001.

Astrocyte mitochondrial morphology was evaluated using the MitoTracker Deep Red FM probe, which accumulates in mitochondria of living cells, allowing their labeling (Figure 5A and B). Automated quantification of mitochondrial morphology indicated that PD-associated mutations decreased average mitochondrial volume per cell (Figure 5C), increased sphericity (Figure 5D), increased branching (Figure 5E), and promoted longer branch length (Figure 5F) compared to control cells. Overall, this demonstrates that the mutations lead to lower mitochondrial presence per cell and more fragmented mitochondria. Additionally, compared to the WT control, *LRRK2* and *PRKN* mutant cells exhibited mitochondria with increased peripheral localization (Figure 5H) and less perinuclear mitochondria (Figure 5I), likely reflecting altered trafficking and transport. This profile was particularly predominant in *PRKN* mutant astrocytes, which also displayed a different pattern from the *LRRK2* mutants. Recently, *PRKN* knockout astrocytes have been reported to exhibit mitochondrial, lysosomal, and proliferation dysfunction (Goldman et al., 2025), which is compatible with these results. A previous study has also demonstrated reduced GFAP reactivity in astrocytes from *PRKN*-mutated human brains and organoids (Kano et al., 2020); as a component of the cytoskeleton, this may explain changes in mitochondrial cellular location and morphology.

**Figure 5.**
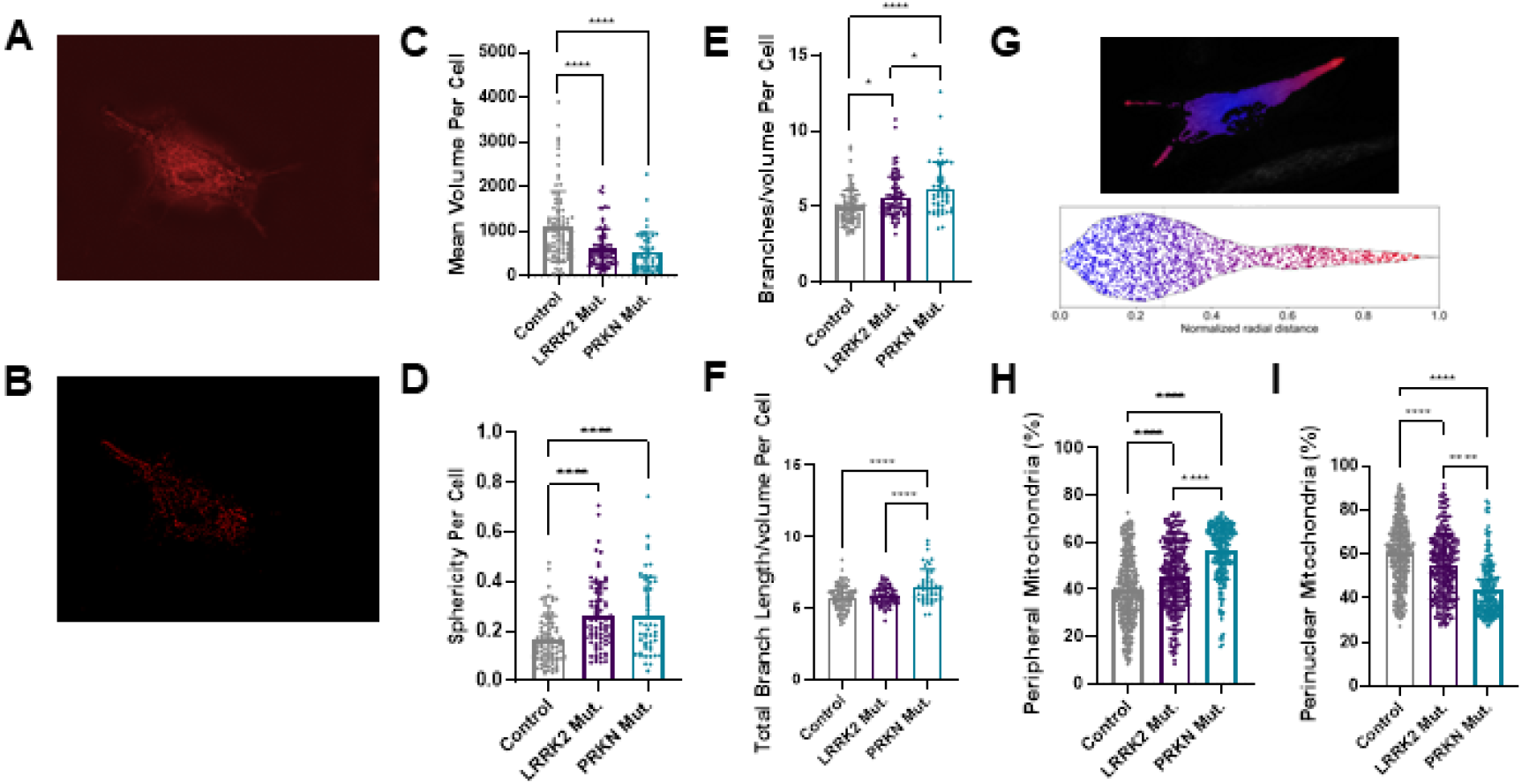
Mitochondria are more fragmented and peripheral in *LRRK2* and *PRKN* mutants. **A)** Representative microscopy with the MitoTracker Deep Red FM probe. **B)** Representative deconvolved image of mitochondria. **C)** Average quantified volume of mitochondria per cell. **D)** Mitochondrial sphericity per cell. **E)** Branching per mitochondrial volume per cell. **F)** Total branch length per mitochondrial volume per cell. **G)** Representative microscopy of a cell and its respective radial mitochondrial location estimate, from the most perinuclear (in blue) to the most peripheral (in red). **H)** Representative plot of the mitochondrial location where each point refers to a mitochondrion in each cell. **I)** Percentage of peripheral mitochondria. Graphs represent Means ± SD. Symbols show individual cell data. One-way ANOVA *P_adj_ < 0.050; **P_adj_ < 0.010; ***P_adj_ < 0.001; ****P_adj_ < 0.0001.

Given these changes in mitochondrial morphology and location, we measured mitochondrial oxygen consumption rates in the astrocytes under physiological growth conditions over the course of 3 days using a Resipher metabolic profile detection system and data platform (Figure 6). We found that control astrocytes displayed marked heterogeneity in oxygen consumption patterns, while *LRRK2* mutants had consistently low oxygen consumption. This pattern differed from that of *PRKN* mutants, where the entire interquartile range was elevated relative to the *LRRK2* mutants. We hypothesize that the higher oxygen consumption in *PRKN* mutants relative to *LRRK2* may be due to loss of mitochondrial oxidative phosphorylation function that is partially compensated for by a defect in mitophagy-mediated mitochondrial clearance. In the literature, such compensatory respiration has been reported in skin fibroblasts and lymphoblasts from individuals with PD and *PRKN* mutations, but not in hiPSC-derived neurons (González-Casacuberta et al., 2019; Castelo Rueda et al., 2023). Neurons and astrocytes display cell-type-specific organelle signatures (Rhoads et al., 2025), including mitochondria, which may impact respiratory differences between these two cell types. In addition, low oxygen consumption has been previously reported in *LRRK2* mutant astrocytes (Ramos-González et al., 2021), a result compatible with our finding in iPSC-derived astrocytes.

**Figure 6.**
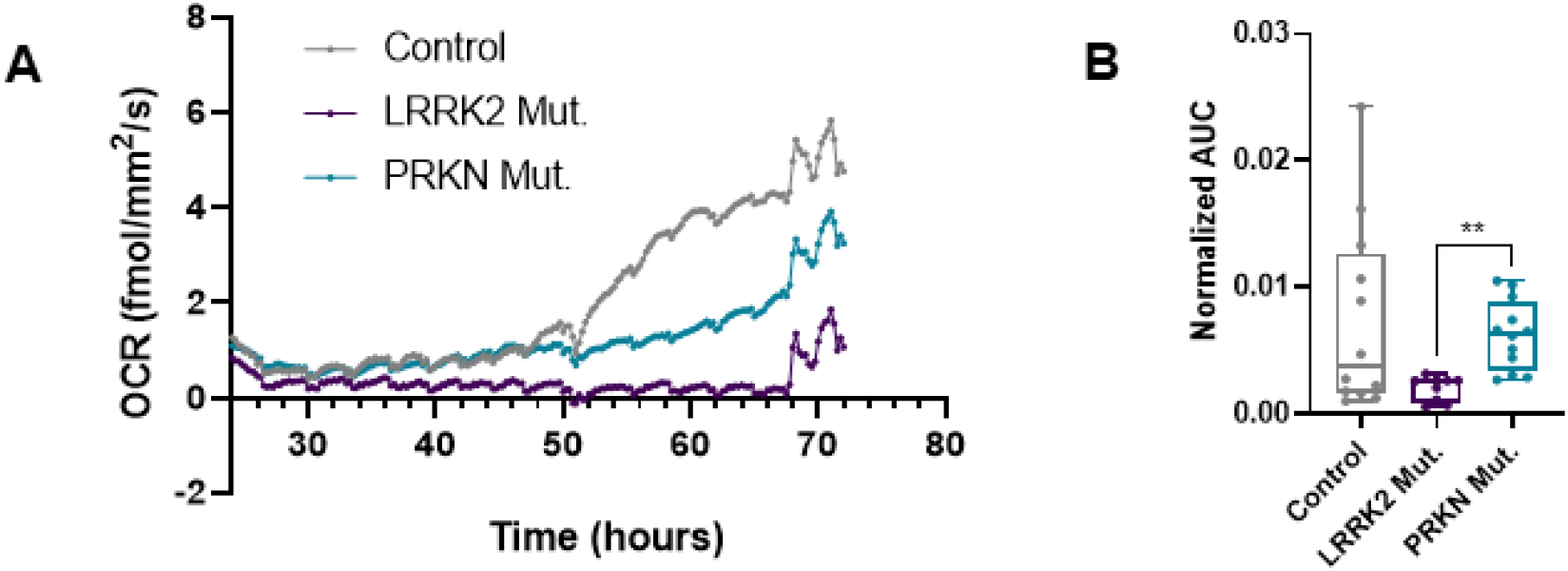
*LRRK2* mutants present decreased oxygen consumption rates. **A)** Representative oxygen consumption traces of control (in gray), *LRRK2* mutant (in purple) and *PRKN* mutant (in turquoise) astrocytes. **B)** Quantifications for the 24-72h period. Data is presented as medians, 10 and 90 percentiles and SDs. Individual symbols are replicates. Kruskal-Wallis with Dunn’s test **P_adj_ < 0.010.

To uncover possible mechanisms involved in the changes in intracellular calcium levels, mitochondrial morphology and metabolic activity, we quantified the relative expression of genes involved in the regulation of mitochondrial morphology and calcium transport (Figure 7). Differences were not statistically significant for any of the analyzed genes, which is not surprising, considering that much of the regulation of these pathways is posttranslational.

**Figure 7.**
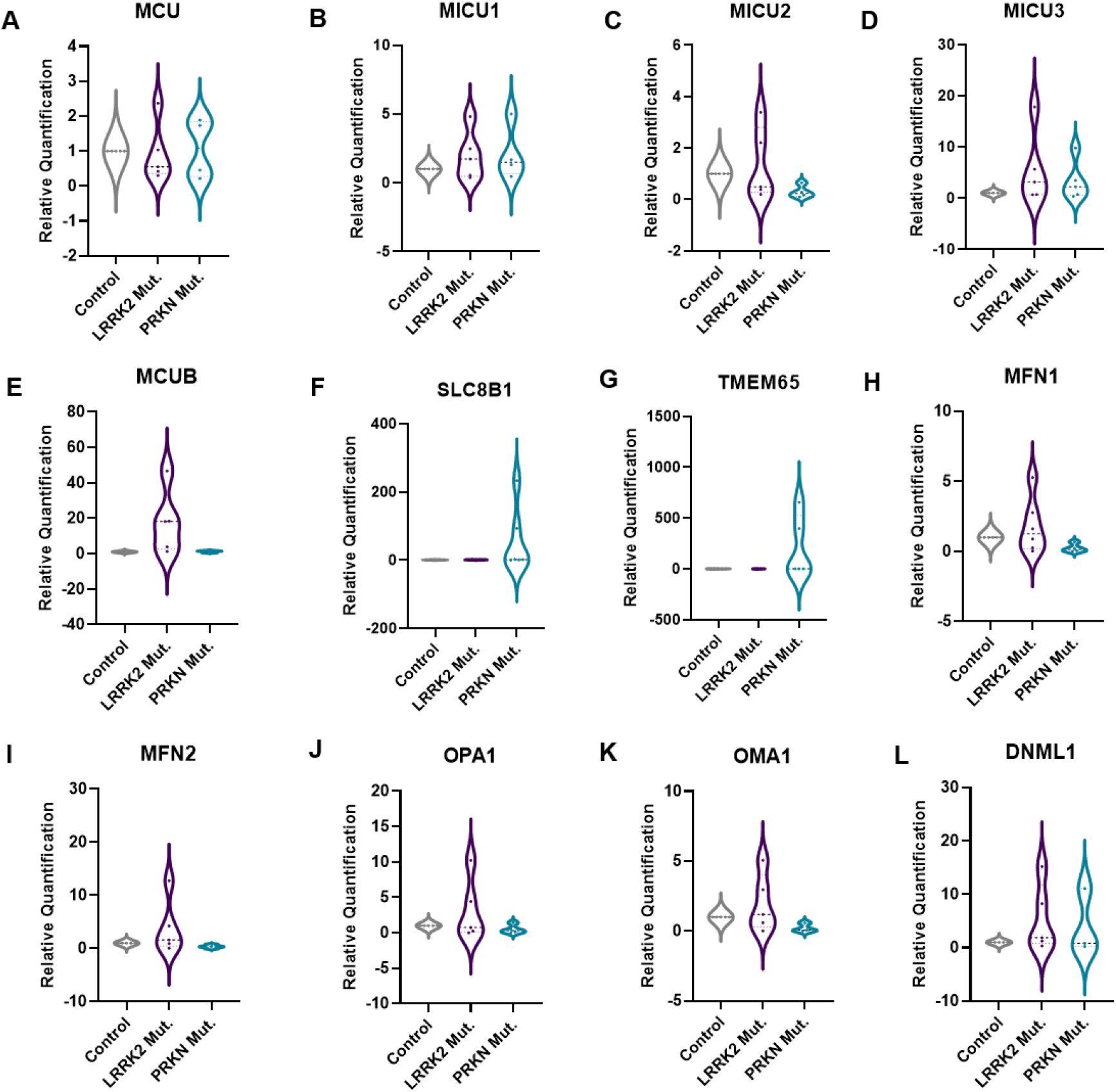
No differences in the gene expression were observed when comparing control (gray) astrocytes, *LRRK2* mutants (purple), and *PRKN* mutants (turquoise). **A-G)** Relative quantification of gene expression for mitochondrial calcium transporters for influx (*MCU*, *MICU1*, *MICU2*, *MICU3* and *MCUB*) and efflux (*SLC8B1*/NCLX and *TMEM65*). **H-L)** Relative quantification of gene expression for mitochondrial fusion (*MFN1*, *MFN2*, *OPA1* and *OMA1*) and fission (*DNML1*/DRP1). One-way ANOVA P>0.05 for all genes.

**Figure 8.**
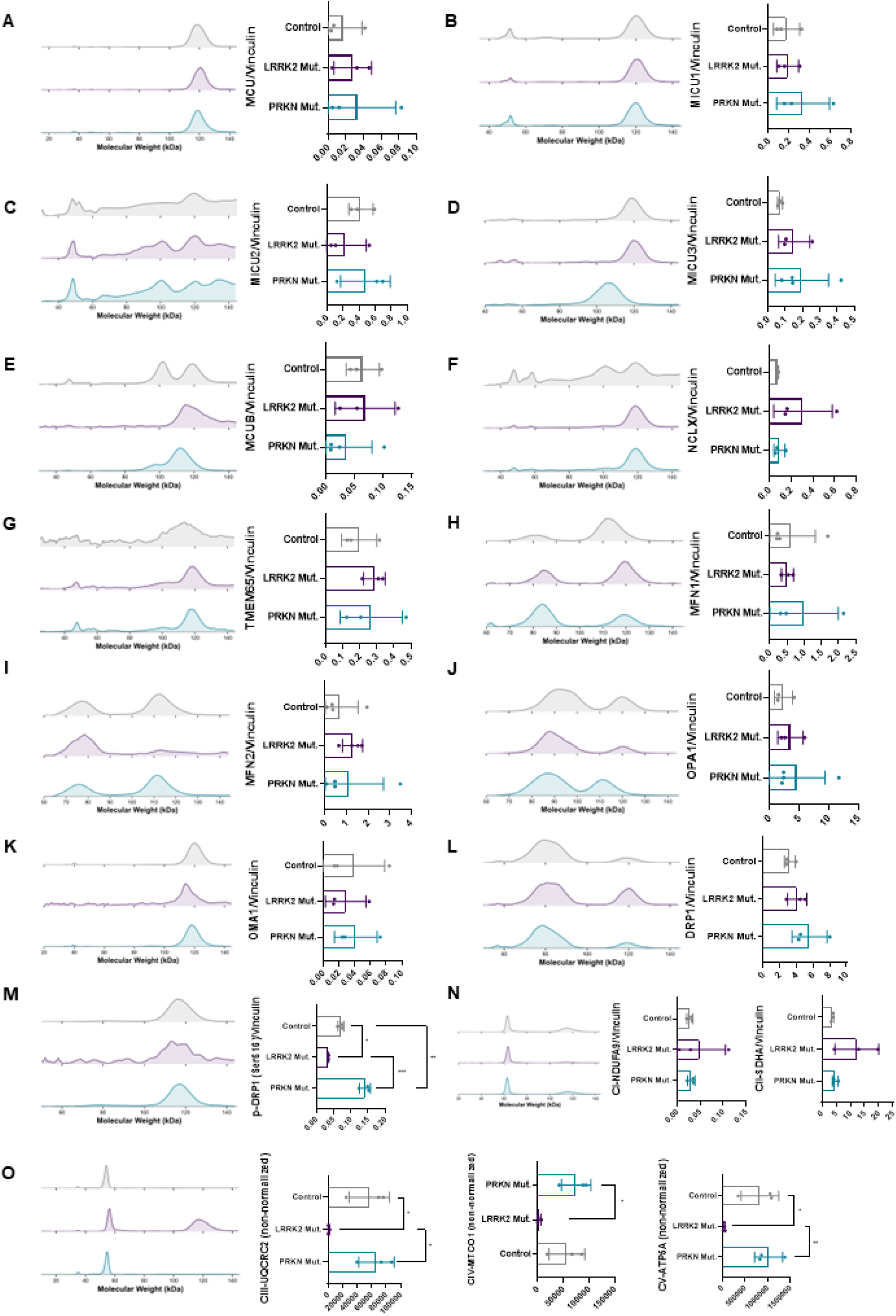
*LRRK2* (purple), and *PRKN* (turquoise) mutations modulate proteins related to mitochondrial morphology and oxygen consumption relative to controls (gray). Representative output electropherograms of capillary electrophoresis westerns are shown on the left of each panel, with one peak for each of the multiplexed proteins (target and housekeeping). **A-G)** Mitochondrial calcium transporter proteins. **H-M)** Mitochondrial dynamics and morphology proteins. **N)** Respiratory complexes I (NDUFA9) and II (SDHA) independent antibodies multiplexed together with Vinculin for normalization. **O)** Respiratory complexes III (UQCRC2), IV (MT-CO1), and V (ATP5A) in an anti-OXPHOS cocktail. One-way ANOVA *P_adj_ < 0.050; **P_adj_ < 0.005; ***P_adj_ < 0.001.

We then analyzed protein content in the cells using a quantitative capillary electrophoresis western detection system adequate for small sample sizes (Jess Automated Western System), and capable of comparing sample and normalization protein detection in the same capillary run. Quantifications of proteins involved in calcium transport in mitochondria did not show significant differences, but DRP1 (a protein involved in mitochondrial fission) Ser616 phosphorylation was significantly altered in the mutant cells, a result which may explain the changes in mitochondrial morphology observed. Indeed, phosphorylation of DRP1 in Ser616 is known to activate mitochondrial fission (Chang et al., 2010), and increased levels of p-DRP1 (Ser616) have been reported in sporadic PD (Santos et al., 2015).

Furthermore, the expression of respiratory complexes II, IV, and V was markedly reduced in *LRRK2* mutants, a finding that strongly correlates with our Resipher results, where the *LRRK2* mutant exhibited a significantly decreased oxygen consumption rate. Similarly, a previous study on iPSC-derived microglia reported that *LRRK2* G2019S mutants had lower oxygen consumption rates compared with controls (Kurniawan et al., 2025), which aligns with findings in mouse embryonic fibroblasts (MEFs) with different *LRRK2* genetic backgrounds (Toyofuku et al., 2019). This result is not universal, as another study on iPSC-derived dopaminergic neurons with a different *LRRK2* mutation (R1441C) showed no oxygen consumption deviation from controls (Williamson et al., 2023). Taken together, these results point to a specific role of the *LRRK2* G2019S mutation in impairing cellular respiration. Indeed, the G2019S mutation in *LRRK2* may affect assembly of respiratory complex IV subunits (Kim et al., 2026), which is compatible with our finding showing highly decreased expression of the complex.

## Conclusions

Using patient-derived iPSCs differentiated into astrocytes, we demonstrate that PD-associated mutations in *LRRK2* and *PRKN* trigger changes in intracellular calcium dynamics, mitochondrial fragmentation, and organelle location reorganization toward the cell periphery, associated with bioenergetic compromise and loss of respiratory complex expression in the *LRRK2* mutant background. By identifying these astrocyte-specific mitochondrial and calcium signatures, we hope our findings highlight the critical role of glial metabolic alterations in the pathophysiology of PD, offering a new perspective on how non-neuronal cells contribute to this neurodegenerative disease.

## Acknowledgements

The authors thank Prof. Alexandre Bruni Cardoso (IQ-USP) for Leica DMi8 microscope access, and Equipamento Multiusuário FSP-USP (FAPESP – Processo 2023/17365-3) for the support with Jess Automated Western experiments. The authors also thank Prof. Júlio Cesar Batista Ferreira (ICB- USP) and Prof. Nadja C. de Souza Pinto (IQ-USP) for kindly donating antibody aliquots.

## Data Availability

The open-source codes developed here were made fully available at https://github.com/cavalcantegc. Raw files are made available on reasonable request.

## Author Contributions

G.C.C. – conceptualization, data curation, formal analysis, investigation, methodology, software, visualization, writing - original draft, writing - review & editing; C.C.S., É.L.V., V.F., and F.G.R. – methodology; P.M.C.A. – supervision, resources, writing - review & editing; A.J.K. – conceptualization, supervision, resources, project administration, funding acquisition, writing - review & editing. All authors read and approved the final manuscript.

